# Morphological and Molecular Phylogenetic Characterization of Three New Marine Goniomonad Species

**DOI:** 10.1101/2024.12.12.627435

**Authors:** Yasinee Phanprasert, Sun Young Kim, Nam Seon Kang, Minseok Jeong, Jong Im Kim, Woongghi Shin, Won Je Lee, Eunsoo Kim

**Author notes:** **Correspondence:** E. Kim, Division of EcoScience, Ewha Womans University, Science Building A114, 52 Ewhayeodae-gil, Seodaemun-gu, Seoul, 03760, South Korea, (or).

## Abstract

Goniomonads are commonly found heterotrophic biflagellates in both marine and freshwater environments. Despite the high genetic diversity inferred from 18S rDNA data, many goniomonad species remain undescribed. In this study, we established a total of 21 marine goniomonad culture strains, and from these, describe three new species by using light microscopy, electron microscopy, and 18S rDNA phylogeny. Molecular sequence analyses suggest the presence of several distinct sub-lineages within the marine goniomonad clade. Two of these are *Goniomonas ulleungensis* sp. nov. and *G. lingua* sp. nov., which are similar in size, flagellar length, appendage & orientation and by having a tongue-like protrusion at the anterior. The two species can be differentiated by the periplast plate pattern with *G. ulleungensis* displaying one additional plate on the right side. *G. duplex* sp. nov. differed from these two species by having two unequal flagella with the longer one trailing posteriorly and having the opposite cell orientation when skidding. Comparative analyses of five marine goniomonad species showed that genetically distinct goniomonad groups can be delineated by morphological data as well, and of several morphological features that are of taxonomic utility, the periplast plate pattern, observable by SEM, is particularly informative in goniomonad taxonomy.

---

Goniomonads, comprising a single genus *Goniomonas*, are colorless biflagellate protists commonly found in both marine and freshwater environments (Lee and Patterson, 2000; Lee et al., 2003; Patterson and Hedley, 1996). In the aquatic food web, they play a role by feeding on bacteria or, alternatively, by serving as food source for zooplankton (Šimek et al., 2023). The genus *Goniomonas* has drawn considerable interest in the study of plastid evolution; some scientists proposed—based on the “Chromalveolate” or “Hacrobia” concepts—that *Goniomonas* originated from photosynthetic ancestors and its plastid was lost secondarily (Cavalier-Smith, 1999; Hackett et al., 2007). However, our more expanded understanding of cryptistan diversity, especially with the identification of additional early diverging, plastid-lacking taxa, points towards the idea that the common ancestor of the Cryptista was most likely plastid-less and *Goniomonas* was likewise ancestrally plastid-lacking. Within the Cryptista, there are multiple sublineages comprising exclusively plastid-lacking members; these include goniomonads (Goniomonadophyceae), katablepharids (Katablepharidophyceae), Palpitia (repesented by a single species *Palpitomonas bilix*; Yabuki et al., 2010). Endohelea (represented by a single species *Microheliella maris*; Yazaki et al., 2022), the Cryptomonad Group 1 (CRY1 or Hemiarmida represeted by *Hemiarma marina*; Shiratori and Ishida, 2016) and CRY3 (recognized by environmental sequence data only with no representative culture strain; Kim and Archibald, 2013). In addition, the nuclear genome of *Goniomonas avonlea* has been sequenced, and contrary to the expectationtion by the “Chromalveolate” or “Hacrobia” concepts, it was found not to carry red algal endosymbiont-derived genes (Cenci et al., 2018; McFadden, 2018), further strengthening the notion that *Goniomonas* was ancestrally non-phosynthetic. Given its close phylogenetic affinity to plastid-bearing cryptophytes, *Goniomonas* serves as a good model for the ancient phagotrophic protist that formed symbiotic association with a red alga and gave rise to photosynthetic cryptophytes (Cenci et al., 2018; McFadden et al., 1994).

So far, four species of *Goniomonas* have been formally described, including *G. truncata, G. pacifica, G. amphinema,* and *G. avonlea* (Kim and Archibald, 2013) with the first three originally described based on light microscopy only without molecular sequence information. *G. truncata—*the type species for the genus—was described from freshwater by Fresenius (1858) as *Monas truncata* that is 7-10 µm long with two flagella about or slightly longer than the cell length. It was described to be oval or rounded at the posterior with a transverse line along the anterior margin; have a flattened body that is slightly curved when viewed from the side; and swim without rotating the body. *G. truncata* was studied by various other researchers, including Mignot (1965) and Ekelund & Patterson (1997), but the surprising 18S rDNA genetic diversity detected among freshwater *G. truncata*-like strains (von der Heyden et al., 2004) suggests the freshwater goniomonads are undersplit and these earlier studies possibly were looking at different goniomonad species. Unfortunately, neither von der Heyden et al. (2004) nor McFadden et al. (1994) described the morphology of their freshwater *Goniomonas* strains when presenting 18S rDNA data. A similar issue exists for two marine *Goniomonas* species: *G. pacifica* and *G. amphinema.* Larsen and Patterson (1990) described the two species based on light microscopic observation. Similar to *G. truncata*, these two colorless marine species have a flattened cell body with a transverse band that runs along the anterior end. More specifically, *G. amphinema* was described to be 5-8 µm long and 4-6 µm wide, obtuse anteriorly and rounded posteriorly. It has two unequal flagella with the longer one, slightly longer than the cell, trails behind and the shorter one, half the cell length, orients anteriorly. *G. pacifica*, on the other hand, was described to be 8-10 µm long and 6-8 µm wide with a truncated anterior end and two equal or sub-equal flagella about ¾ the cell length. One flagellum directs forwards, while the other bends towards the opposite corner of the anterior margin. The cell size ranges for *G. pacifica* and *G. amphinema* have been expanded by the original authors and others in later studies (see summary within Kim and Archibald, 2013). Several 18S rDNA sequences have been acquired and deposited to GenBank under the taxon names *G. pacifica* and *G. amphinema*, but unfortunately most of these do not have records for morphological data. *G. avonlea* was most recently described (Kim and Archibald, 2013) and it differs from *G. truncata* by inhabiting marine environments. It was described to be 8–11 µm long and 6–8 µm wide, similar to the original size description for *G. pacifica*. However, unlike *G. pacifica*, *G. avonlea* has two flagella that are slightly longer than the cell length with one trailing posteriorly. This species is larger than *G. amphinema.* Previous studies based on the phylogenetic analyses of 18S rDNA sequences showed considerable genetic divergence among goniomonad sequences, suggesting the existence of undescribed species (Kim and Archibald, 2013; Martin-Cereceda et al., 2010; von der Heyden et al., 2004).

In this study, we describe three new species of marine goniomonads based on the analyses of 18S rDNA and morphological data: 1) *Goniomonas ulleungensis* sp. nov., 2) *G. lingua* sp. nov., and 3) *G. duplex* sp. nov. Our phylogenetic analyses revealed that the marine goniomonads are classified into several genetically distinct groups despite their morphological conservation detectable under a light microscope. Nonetheless, we demonstrate that cell size and the flagella length and orientation as well as the ultrastructure features, particularly the periplast plate pattern, are informative and can be used in delineating marine goniomonad species.

## MATERIALS AND METHODS

### Sample collection and culturing

Seawater samples were collected from 17 coastal locations in South Korea between 2017 and 2023 and one location within Prince Edward Island of Canada in 2011 (Table 1). Field samples were enriched in a modified ESM medium (Okaichi et al., 1982; addition of 5ml Seawater-802 [ATCC medium 1525] medium in 1-L volume), AF-6 medium (Andersen et al., 2005) with barley grains, or SL medium (= 1% v/v LB in sterile seawater). Goniomonad cells were isolated by a single-cell isolation with a finely drawn Pasteur pipette or by a serial dilution method. A total of 21 clonal goniomonad cultures, alongside uncharacterized bacteria from the field seawater samples, were established and maintained between 17°C and 21°C under dark or with a 12-hour light cycle.

**Table 1.** Collection date, location, and GenBank ID for the isolates obtained and analyzed in this study. An asterisk symbol denotes the strains that remain—as of October 2024—alive and regularly transferred to fresh medium. Abbreviation: MPRBK, Marine Protist Resource Bank of Korea.

| Strains<br>(culture ID) | Date | Location (latitude-longitude) | GenBank ID<br>(18S rDNA) |
| --- | --- | --- | --- |
| *Goniomonas ulleungensis* |  |  |  |
| UH_01*<br>(MPRBK-0001) | 2023.10.18 | Hyeonpo Port, Ulleung-do, South Korea<br>(37.5272, 130.8296) | PQ432910,<br>PQ436815 |
| JY_01 | 2023.06.27 | Nonjinmul, Yerae-dong, Seogwipo-si, Jeju-do, South Korea<br>(33.2370, 126.3899) | PQ436665 |
| KM017* | 2022.04.21 | Seonyu Island, Gunsan, Jeollabuk-do, South Korea<br>(35.8137, 126.4156) | PQ490427 |
| KM022* | 2022.05.24 | Seodo beach, Yeosu, Jeollanam-do, South Korea<br>(34.0553, 127.2946) | PQ443870 |
| KM066 | 2021.05.06 | Seodo Harbour, Yeosu, Jeollanam-do, South Korea<br>(34.0547, 127.2979) | PQ443871 |
| KM071* | 2022.07.26 | Bongpyong beach, Uljin-gun, Gyeongsangbuk-do, South Korea<br>(37.0436, 129.4144) | PQ443872 |
| KM073* | 2022.06.30 | Dugok beach, Namhae-gum, Gyeongsangnam-do, South Korea<br>(34.7684, 127.9076) | PQ443954 |
| KM074* | 2022.06.29 | Gorageum beach, Goheung-gun, Jeollanam-do, South Korea<br>(34.4779, 127.1152) | PQ443955 |
| KM097* | 2022.04.02 | Gwangam beach, Changwon, Gyeongsangnam-do, South Korea<br>(35.1025, 128.5000) | PQ443957 |
| SambongG17 | 2017.12.02 | Sambong beach, Taean-gun, Chungcheongnam-do, South Korea<br>(36.3721, 127.3496) | PQ444059 |
| *Goniomonas lingua* |  |  |  |
| GW*<br>(MPRBK-0003) | 2011.08.27 | Greenwich beach, Prince Edward Island, Canada<br>(46.4503, -62.7125) | PQ448234,<br>PQ448237 |
| KM021* | 2022.04.21 | Gotji beach, Taean-gun, Chungcheongnam-do, South Korea<br>(36.4878, 126.3333) | PQ443867 |
| GamamiA4 | 2017.11.24 | Gamami beach, Yeonggwang-gun, Jeollanam-do, South Korea<br>(35.3997, 126.4091) | PQ444058 |
| ManseongA5 | 2018.01.12 | Manseongri beach, Yeosu-si, Jeollanam-do, South Korea<br>(34.7769, 127.7451) | PQ444054 |
| *Goniomonas duplex* (and related strains) |  |  |  |
| PT_04*<br>(MPRBK-0002) | 2023.10.31 | Padori beach, Taean-gun, Chungcheongnam-do, South Korea<br>(36.7440, 126.1324) | PQ436667,<br>PQ436816 |
| JG2_01 | 2023.11.07 | Gujwa-eup, Jeju-do, South Korea<br>(33.5433, 126.8288) | PQ436670 |
| BatgaeB2 | 2017.12.02 | Batgae Beach, Taean-gun, Chungcheongnam-do, South Korea<br>(36.5308, 126.3307) | PQ444052 |
| ManseongB2 | 2018.01.12 | Manseongri beach, Yeosu-si, Jeollanam-do, South Korea<br>(34.7769, 127.7450) | PQ444035 |
| ManseongC7 | 2018.01.12 | Manseongri beach, Yeosu-si, Jeollanam-do, South Korea<br>(34.7769, 127.7450) | PQ444034 |
| PoongryuA1 | 2018.01.12 | Poongryu beach, Goheung-gun, Jeollanam-do, South Korea<br>(34.6642, 127.2295) | PQ444042 |
| *Goniomonas avonlea* |  |  |  |
| PoongryuC10 | 2018.01.12 | Poongryu beach, Goheung-gun, Jeollanam-do, South Korea<br>(34.6642, 127.2294) | PQ444071 |

### DNA preparation, sequencing, and phylogenetic analyses

From each 10-20-ml clonal culture, cells were collected by centrifugation at 2,000-9,500 x *g* for 2-5 min or by gentle vacuum filtration onto a 0.8 µm polycarbonate filter. The total DNA was extracted from a cell pellet using a DNeasy Blood & Tissue kit (QIAGEN, Hilden, Germany) or a Exgene Cell SV kit (GeneAll Biotechnology, South Korea) following the manufacturer’s recommended protocols. Polymerase chain reaction (PCR) was performed to amplify 18S rDNA regions using “universal” eukaryotic primers (Table S1). The amplified products were cleaned and processed for Sanger sequencing (Applied Biosystems 3730xl DNA analyzer) at Cosmogenetech co, Ltd (Seoul, South Korea). Output reads were inspected for corrections, and, if needed, merged using 4Peaks (v. 1.8; by A. Griekspoor and T. Groothuis, nucleobytes.com), Mesquite ver. 3.81 (Maddison and Maddison, 2023), Geneious (v. 8.15; Kearse et al., 2012), or Genetic Data Environment program v. 2.6 (Smith et al., 1994). For a representative strain for each of three new species described in this study (i.e. the strains UH_01, GW, and PT_04), two independent PCR and sequencing experiments were conducted to ensure the accuracy of sequence data. Newly obtained 18S rDNA sequences have been deposited in GenBank with the accession numbers as shown in Table 1.

The newly acquired 18S rDNA sequences were manually aligned with reference sequences obtained from GenBank using Mesquite (ver. 3.81; Maddison and Maddison, 2023). Poorly aligned regions were identified and were marked for exclusion, yielding the final alignment (Data S1) with the total number of 1,901 characters (after trimming) and 107 sequences. The maximum likelihood analyses were performed using IQ-TREE (ver. 2; Minh et al., 2020) with the TN+F+I+G4 model identified by ModelFinder (Kalyaanamoorthy et al., 2017). Maximum parsimony analyses were conducted using MPBoot tool available on the CIPRES Science Gateway (XSEDE) (Miller and Pfeiffer, 2010). Bootstrap analyses were based on 1,000 resamples. Figtree (ver. 1.1.4; Rambaut, 2018) was used to visualize the trees.

### Light microscopic observation

Living goniomonad strains were observed with the differential interference contrast (DIC) optic under a Nikon ECLIPSE Ti2-E inverted microscope equipped with a Nikon digital sight 10 camera (image acquisition using NIS-Elements BR); a Leica DMR microscope (Germany) equipped with a Zeiss Axiocam HR digital camera (image acquisition using Axiovision 4.6); or a Zeiss Axio Imager A2 microscope (Germany) equipped with a Zeiss AxioCam 712 color photomicrographic system. Acquired images were subjected to cell size measurements using the measurement feature of Adobe Photoshop 2022. Boxplots were generated to visualize cell size data using R, utilizing the geom_boxplot() function from the ggplot2 package (R Core Team, 2022; Wickham, 2016).

### Scanning electron microscopy (SEM)

For scanning electron microscopy, drops of cultured cells were placed onto a poly-L-lysine-coated German glass coverslip. Following the protocol by Lee and Simpson (2014), protist cells were first vapor fixed with 4% (w/v) OsO_4_ for 30 min at room temperature. Subsequently, a drop of 4% (w/v) OsO_4_ was directly added to the vapor fixed sample at the final concentration of ∼1% (w/v). The cells were then rinsed with sterile distilled water, dehydrated using a graded ethanol series [3 min each at 30%, 50%, 70%, 80%, 90%, 95% (2 changes), and 100% (3 changes)], and critical point dried with CO_2_ (EM CPD300, Leica, Wetzlar, Germany). The filters were mounted on stubs, sputter-coated with gold (MC1000, Hitachi, Tokyo, Japan), and examined using a high-resolution Zeiss Sigma 500 VP FE-SEM (Carl Zeiss).

### Transmission electron microscopy (TEM)

For each strain, cells were harvested from 100 to 500-ml culture material by centrifugation at 3,000 x *g* for 5-10 min. After decanting the supernatant, the cells were fixed in 2.5% glutaraldehyde solution in marPHEM buffer (600 mM, pH 7.4) (Montanaro et al., 2016). After washed in marPHEM buffer, specimens were post-fixed with 1% osmium tetroxide for 1 hour, dehydrated with ethanol, and embedded in epoxy-resin. Ultrathin sections, approximately 60-70nm in thickness, were cut by an ultramicrotome (Leica EMUC7) using a diamond knife. Sections were contrasted with uranyl acetate followed by lead citrate and observed with H-7650 TEM (Hitachi, Tokyo, Japan) at an accelerating voltage of 80 kV.

For whole-mount TEM, a dense or 10-20x concentrated culture material was used with or without fixation in glutaraldehyde (2.5% final concentration). Cells were concentrated by centrifuging ∼1ml culture at 3,000 x *g* for 5 min, followed by carefully removing the supernatant. A small drop of cells (5 µl) was placed onto a coated grid Formvar-carbon coated EM grids. The grids were stained with 1-2% uranyl acetate and the liquid was removed using filter paper. The grids were observed using a H-7650 transmission electron microscope (Hitachi, Tokyo, Japan) at a voltage of 80 kV.

## RESULTS

In this work, we isolated a total of 21 marine goniomonad strains, all from coastal water samples in South Korea except for one that was from a sandy beach, Prince Edward Island, Canada. From these strains, we describe three new goniomonad species using a combination of light microscopy, SEM, TEM (standard and whole-mount), and 18S rDNA phylogeny.

Note that in describing the goniomonad cell morphology, Kim and Archibald (2013) defined the ventral side as the side that is faced towards the surface on which the cells move along, and the dorsal side is the oppostie to the ventral side. However, our study revealed that some goniomonad sub-lineages switched the orientation of its body when skidding, so with this definition, it becomes a bit confusing to compare different goniomonad species Therefore, we follow the terminology proposed by Mignot (1965) in that the narrow side where the flagellar emerge is defined as the dorsal surface and the opposite side as the ventral surface. The two wide sides then are defined as the right or left sides.

### Light microscopy

All marine goniomonad strains investigated in this study exhibited gross morphological similarities with the flattened cell body rounded (or pointed) at the posterior and truncated or obstruse at the anterior (Figures 1, S1). Cells were colorless with a conspicuous transverse band of ejectisomes running along the anterior margin. Two flagellar arose from an anterior depression. The newly acquired strains were maintained with co-cultured bacteria that likely came from the same water source; goniomonad cells presumably feed on at least some of these through its opening along the anterior margin. Cells rotated in the water column and skidded along the surface while rapidly changing the direction of its move (Movies S1-S3).

**FIGURE 1.**
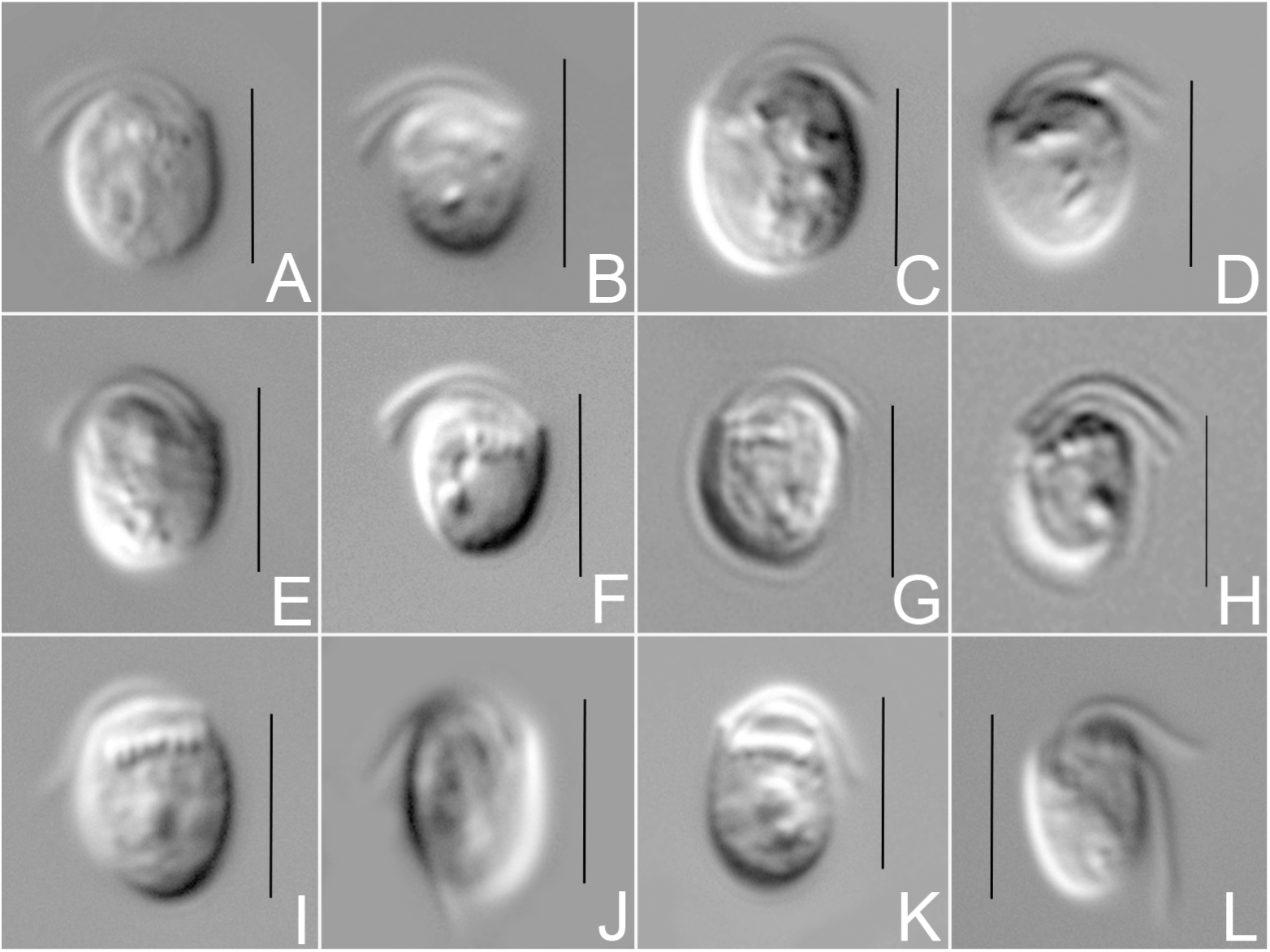
Differential interference contrast (DIC) light microscope images of three marine goniomonad species. Cells were compressed with two flagella emerging from an anterior depression. A row of large ejectisomes ran across the cell’s anterior margin. **A-D**. *Goniomonas ullengensis* strain UH_01. **E-H**. *G. lingua* strain GW. **I-L**. *G. duplex* strain PT_04. Unlike UH_01 and GW, which possessed two nearly equal length flagella, the strain PT_04 displayed two unequal flagella with the longer one extending along the anterior margin and then bending towards the posterior end. Scale bars = 5 μm.

Four strains of *Goniomonas ulleungensis—*UH_01, KM071, KM097, SambongG17— were subjected to light microscopic imaging and cell size measurements (Figures 1, 2; Data S2). Of these, three—UH_01, KM097, SambongG17—had similar size ranges with the cell length between 4-5 µm and the width between 3.3-4.3 µm. KM071, despite having >99% 18S rDNA sequence similarity with UH_01, was smaller with the average length of 3.22 ± 0.51(SD, n=26) µm and the average width of 3.49 ± 0.42 (SD) µm. KM071 also differed from other strains in that it is stretched a bit out laterally with the average length-to-width ratio of 0.93, whereas other were elongated vertically (Figures 1A-D, S1A-F). Two almost equal-length flagella that are slightly longer than the cell arose from an anterior depression. The movement pattern was observed from the strain UH_01. Many cells were in the water column; it tumbled and did not move in a straight line. Cells on the bottom of the culture flask were mostly sedentary with both flagella oriented anterior laterally (Movie S1). When the cell skidded, its left side faced towards the surface (Movie S1).

**FIGURE 2.**
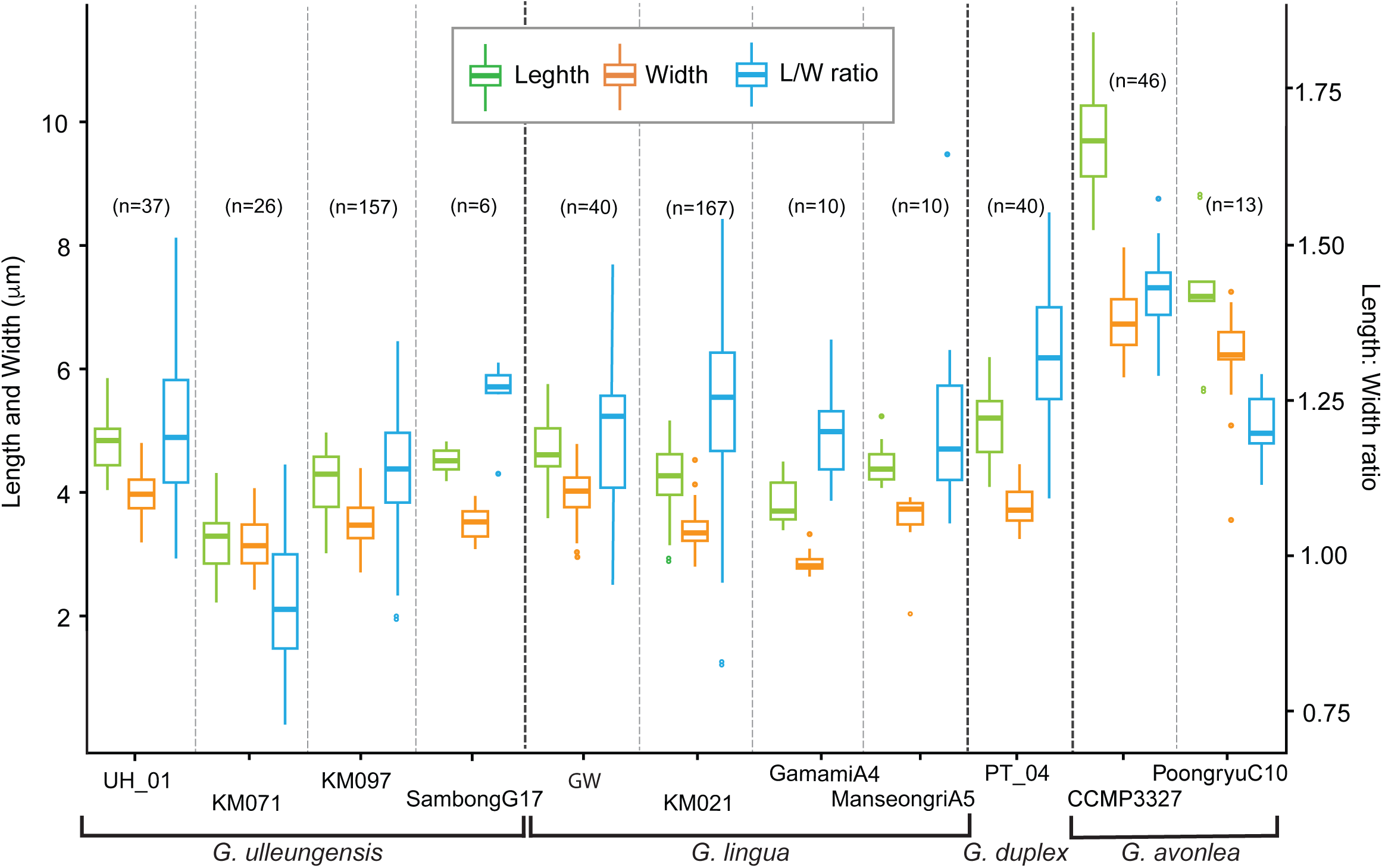
Boxplots displaying cell size measurement data for eleven marine goniomonad strains. The cell length and width as well as the length/width ratio are shown with the vertical lines representing the average values.

Light microscopic imaging data for *Goniomonas lingua* were acquired from the strains GW, KM021, GamamiA4, and ManseongriA5 (Figures 1E-H, S1). All strains exhibited a slightly elongated body shape with the length-to-width ratio around 1.2. Cells were 3.8-5µm long and 3-4 µm wide with the strain GamamiA4 being slightly smaller than the others. Two almost equal-length flagella of slightly longer than the cell arose from an anterior depression. The cell movement for *G. linguia* was characterized by observing the strain GW (Movie S2). The pattern of swimming and skidding was very similar between *G. lingua* and *G. ulleungensis*. For both, cells were mostly in the water column and skidding cells were not as numerous when compared to *G. avonlea* CCMP 3327 (Kim and Archibald, 2013) or *G. duplex* PT_04 (Movie S3). Further, both species had the same cell orientation when skidding, with its left side facing the surface and the right side facing upwards.

*Goniomonas duplex* strain PT_04 (Figure 1I-L) was observed to be 5.10 ± 0.57 (SD, n=40) µm long and 3.83 ± 0.31 (SD) µm wide, with the average length-to-width ratio of 1.33. Cells had two unequal flagella with the anterior flagellum (AF) about the cell length and the longer posterior flagellum (PF). When skidding on the surface, the PF was curved towards the posterior end (Movie S3). Like *G. avonlea* CCMP 3327, this species had its right side facing the surface, which is opposite to *G. ulleungensis* and *G. lingua* (Movie S3). Further, like *G. avonlea, G. duplex* exhibited the behavior of lifting its posterior end while feeding a bacterium on the surface through its “mouth” (Movie S3). Such feeding behavior was not observed for *G. ulleungensis* or *G. lingua*.

A new *Goniomonas avonlea* strain acquired in this study (PoongryuC10) was 7.27 ± 1.12 (SD, n=13) µm long and 6.03 ± 0.92 (SD) µm wide with the average length-to-width ratio of 1.21.

### Scanning electron microscopy

SEM data were acquired from three strains of *Goniomonas ulleungensis* (UH_01, KM071, KM097), two strains of *G. lingua* (GW, KM021), and the *G. duplex* strain PT_04 (Figures 3-5, S2). Examination of multiple different cells from each strain suggests the periplast plate patterns are well conserved within each species and can be used as a taxonomic marker.

**FIGURE 3.**
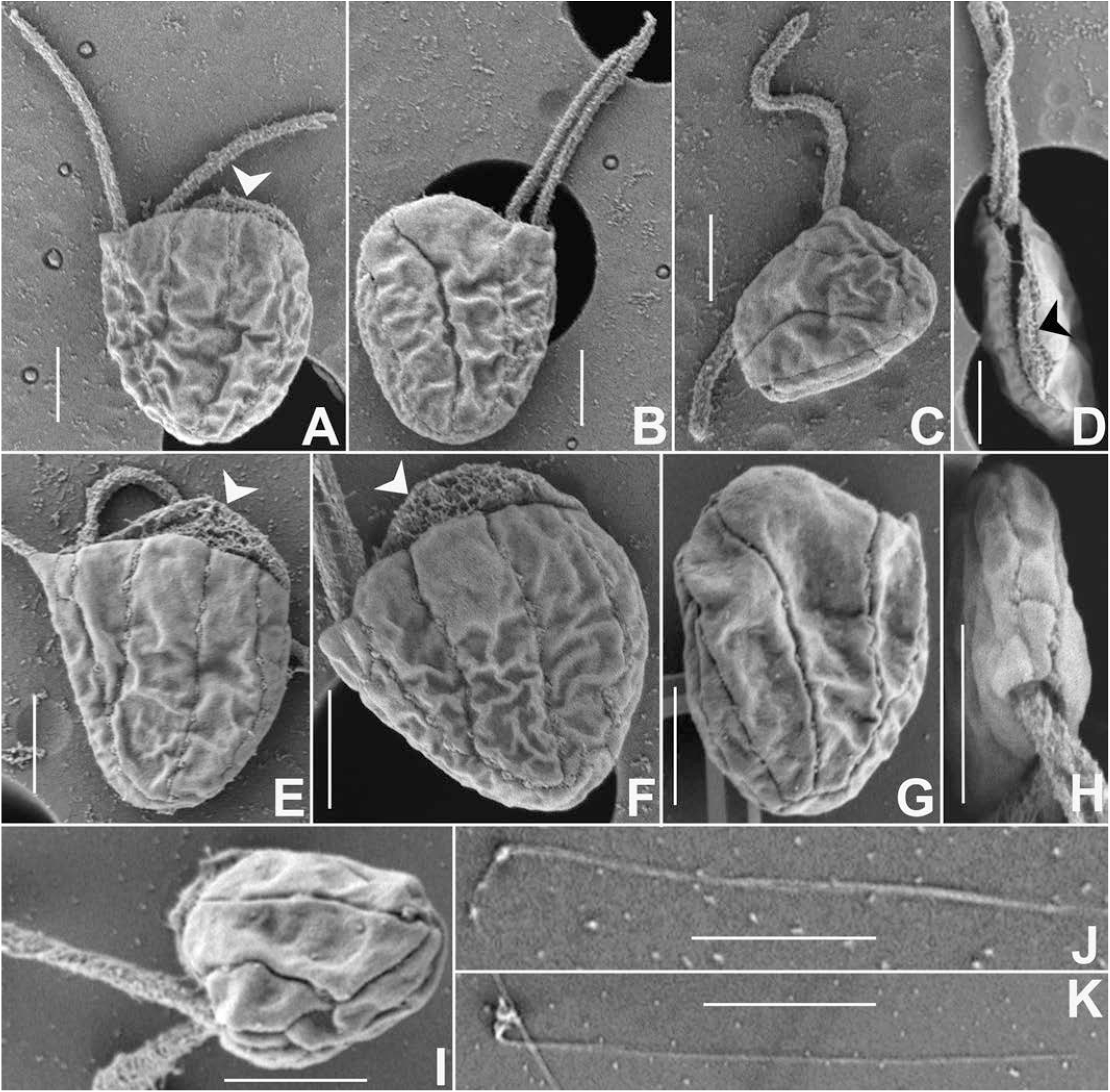
Scanning electron microscope (SEM) images for *Goniomonas ullengensis* strain UH_01 showing the right (**A, E, F**), and left (**B, G**) sides. The right side displayed four periplast plates (**E, F**) whereas the left side had three with the middle one protruding anteriorly with a rough textured pad affixed towards the cell’s interior (arrowheads in **A, D, E, F**). A cell was compressed (**C, D**) with the dorso-ventral sides supported by a narrow periplast strip or two (**C, H, I**). An unfurled small ejectisome was typically longer than the cell length and consisted of two parts with one much shorter than the other (**J, K**). Scale bars = 1 µm.

*Goniomonas ulleungensis* and *G. lingua* displayed a tongue-like structure at the cell anterior (arrowheads in Figures 3A, 4A). For both species, about 2/3 of the anterior margin in the left side extended upwards, creating the obtuse cell morphology, visible from the right side (Figure 3A). The protruded region had an interior padding with rough surface texture (Figure 3D). In *G. ulleungensis*, four periplast plates were observed on the right side and three periplast plates on the left side. Periplast strips supporting the dorsal and ventral sides could be partially seen from the right and left side views (Figure 3A, 3B). The small ejectisomes were expelled into a thin bipartite rod that was typically longer than the cell length (Figure 3J, 3K).

**FIGURE 4.**
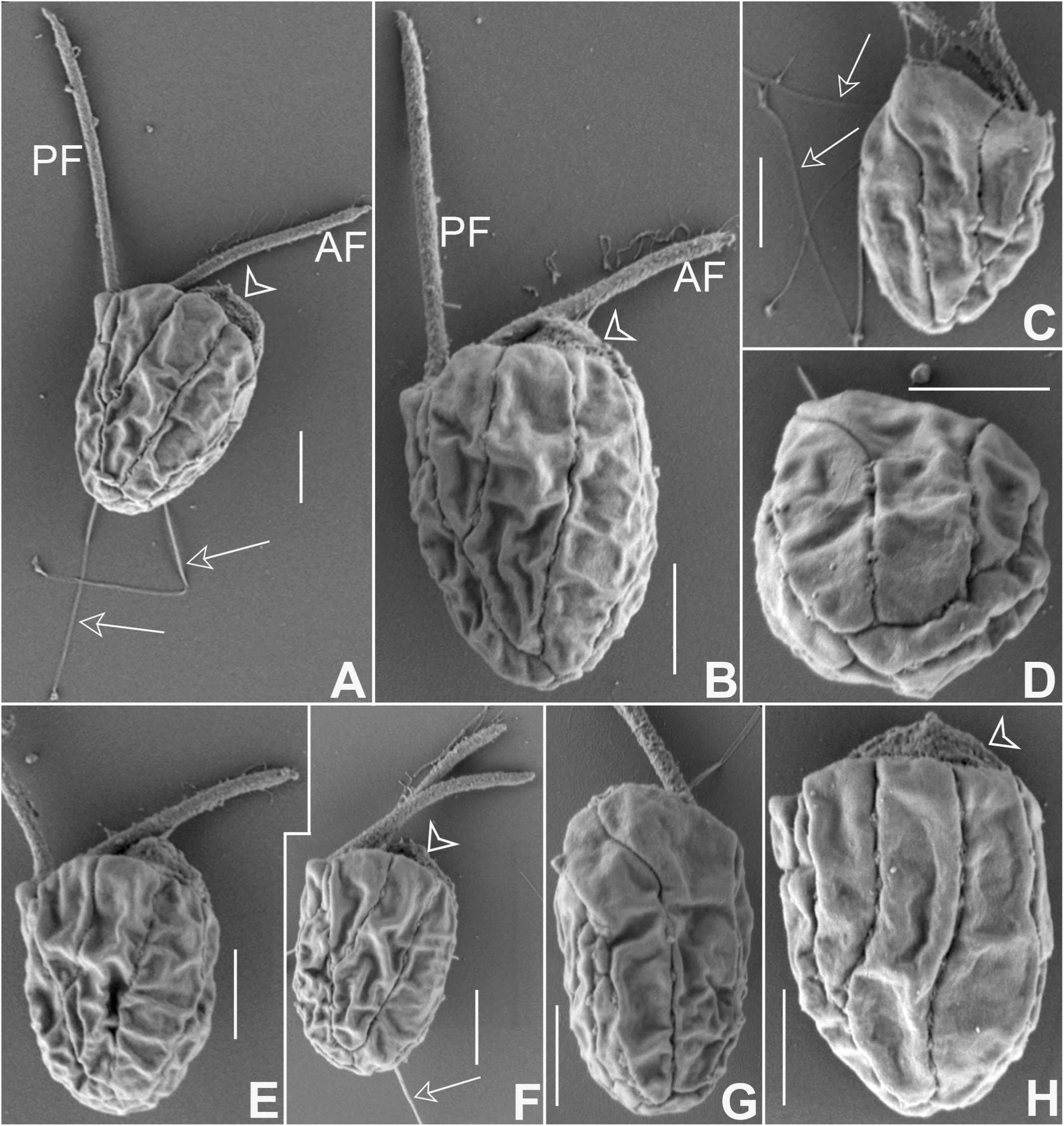
Scanning electron microscope (SEM) images for *Goniomonas lingua* strain GW showing the right (**A, B, E, F, H**), and the left (**C**) sides. Both sides displayed three periplast plates. Like UH_01, the middle periplast of the left side is protruding anteriorly (arrowheads in **A, B, F, H**). Two near equal-length flagella arose from an anterior side with the anterior flagellum (AF) bearing hairs (**A, B**). Arrows (**A, C, F**) indicate small ejectisomes, which consist of two parts with shorter one expelled first from the cell (**A**). The dorsal and ventral sides are supported by a narrow periplast strip or two (**D, G**). Scale bars = 1 µm.

*Goniomonas lingua* had a similar periplast plate arrangement to that for *G. ulleungensis* with three periplast plates on the left side. In both *G. ulleungensis* and *G. lingua*, the upper area of the middle periplast plate in the left side was fan shaped (Figures 3B, 4C), which formed their characteristic tongue-like structure (arrowheads in Figure 4A, 4B, 4F, 4H) when viewed from the right side. However, *G. lingua* displayed three periplast plates on the right side; the far-right periplast plate on the right side of *G. ulleungensis* was absent (or reduced and shifted into the ventral side) in *G. lingua* (Figures 4A, S2C). The discharged small ejectisomes were slightly longer than the cell length (Figure 4C) and had a bipartite structure consisting of the shorter bird beak-like part and the longer straight part.

*Goniomonas duplex* had a more elongated cell body compared to *G. ulleungensis* and *G. lingua*. Two flagella were unequal with the shorter, hairy anterior flagellum (AF) and the longer, smooth posterior flagellum (PF) (Figure 5B). Like *G. ulleungensis* and *G. lingua*, *G. duplex* discharged the bipartite small ejectisomes (Figure 5C). It had three periplast plates on the right side and five on the left side (Figure 5A, 5B). The tongue-like structure observed in *G. ulleungensis* and *G. lingua* was absent in *G. duplex*.

**FIGURE 5.**
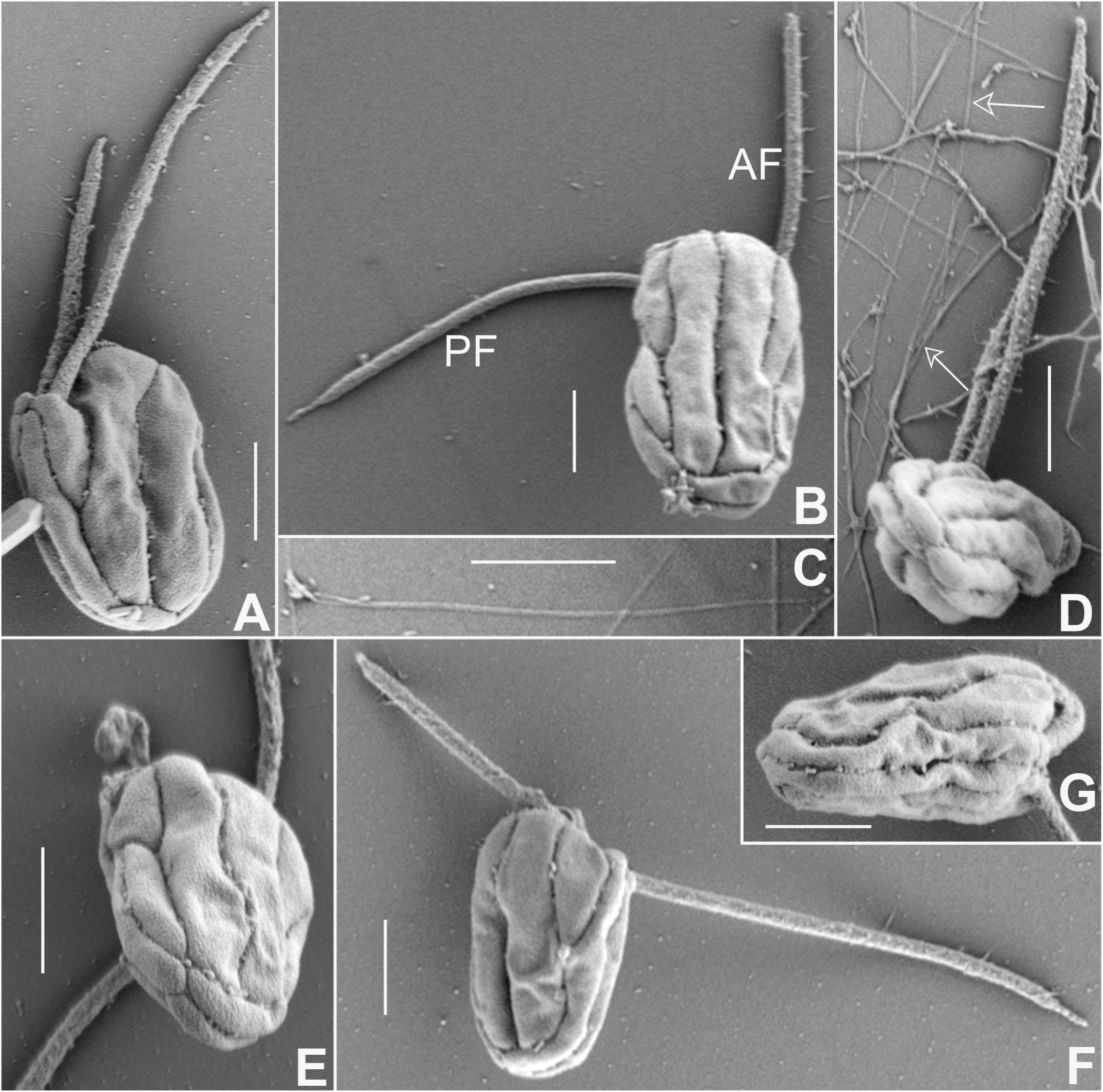
Scanning electron microscope (SEM) images for *Goniomonas duplex* strain PT_04. The left (**A**) and right (**B**) side views, showing three periplast plates on the left side and five on the right side. The anterior flagellum (AF) was about the cell length and bears hairs whereas the posterior flagellum (PF) was smooth and ∼1.5x the cell length (**A, B, F**). The small ejectisomes expelled during the specimen preparation were observed (**C,** arrow in **D**). The dorsal and ventral sides were supported by two narrow periplast plates (**D, E, F, G**). Scale bars = 1 µm.

### Transmission electron microscopy

The whole-mount TEM was helpful in describing the flagellar appendages and the discharged ejectisomes (Figures 6-8). Skipping the cell fixation in glutaraldehyde facilitated the firing of ejectisomes while glutaraldehyde fixation was necessary to conserve the morphology of protist cells. Three strains—*Goniomonas ulleungensis* UH_01, *G. lingua* GW, and *G. duplex* PT_04— were subjected to whole-mount TEM. All three strains were similar in that one flagellum had hairs on whereas the other was smooth. In case for *G. ulleungensis* and *G. lingua*, the flagella hairs were up to several µm long (Figures 6A, 7A). For all three strains, the discharged small ejectisomes were observed with their length typically longer than the cell length (Figures 6C, 6D, 7B, 7F, 8C, 8D). The small ejectisomes had two parts with the shorter part attached to the longer part at a 90-degree angle or greater. The discharged large ejectisomes were observed for *G. ulleungensis* UH_01 (arrowheads in Figure 6C); they were 8–10 µm long and 0.8–1 µm wide. Similar to *G. avonlea* (Kim and Archibald, 2013), the large ejectisomes of *G. ulleungensis* had an outer mesh-like layer over the inner rod. An elongated rod structure with a globular end was observed for *G. lingua* GW (Figure 7D, 7E); it was 3–6µm long and 0.3 µm wide. This might be the large ejectisomes that were partially discharged without the outer mesh-like layer, but more studies are needed to confirm this. We were not able to observe the discharged large ejectisomes for *G. duplex* PT_04 under the preparation conditions used in this study.

**FIGURE 6.**
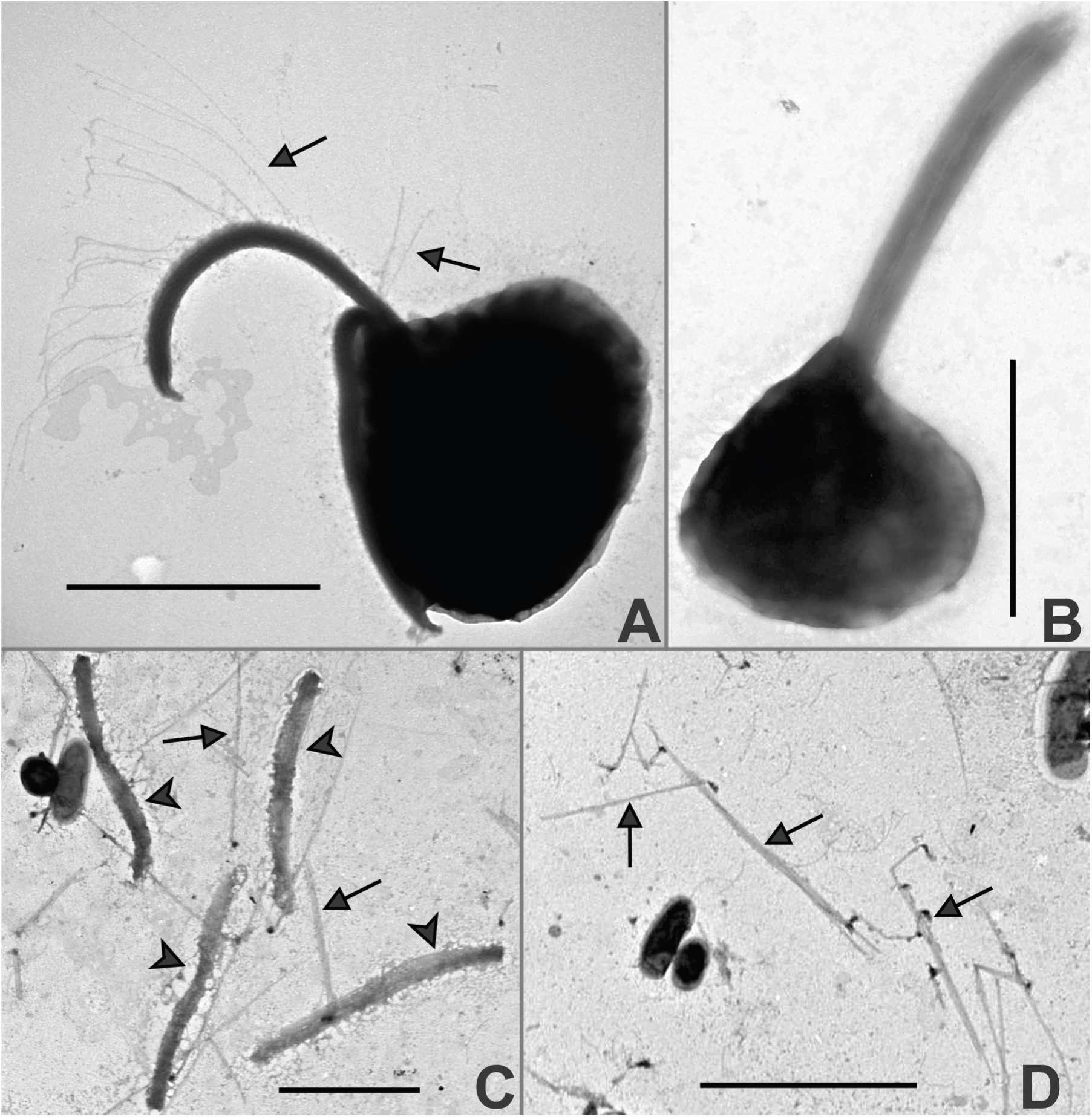
Whole-mount transmission electron microscope (TEM) images from *Goniomonas ullengensis* strain UH_01. Glutaraldehyde fixation preserved the protist cell morphology (**A, B**) whereas no fixation facilitated the expulsion of ejectisomes from the cells while drying on a metal grid (**C, D**). Two flagella were almost equal length (**B**), but only one of them had hairs (arrows, **A**). Both the large (arrowheads, **C**) and small (arrows; **C, D**) ejectisomes were longer than the cell length when fully unfurled. Scale bars = 5 μm.

**FIGURE 7.**
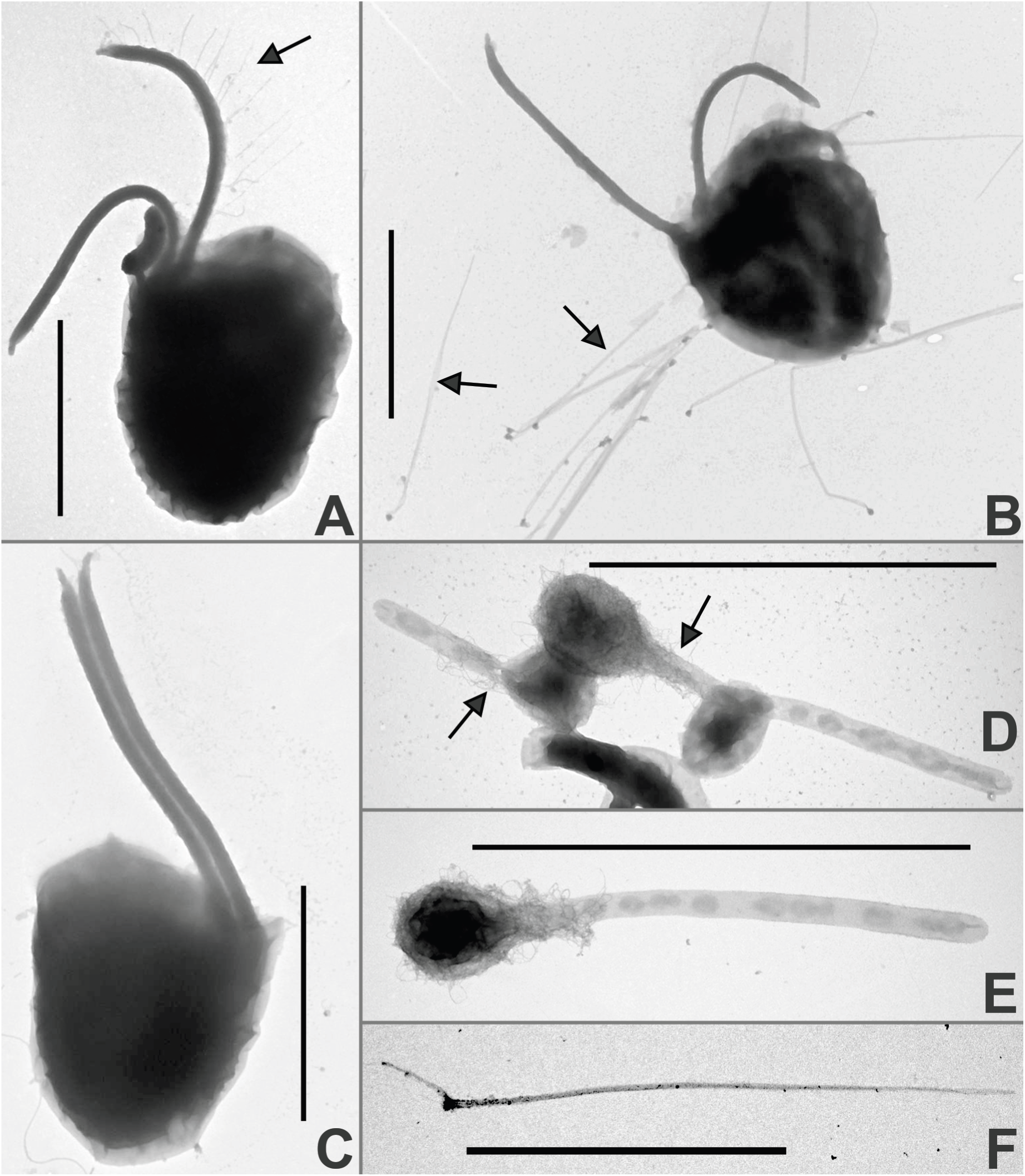
Whole-mount transmission electron microscope (TEM) images from *Goniomonas lingua* strain GW. The protist possessed two flagella of almost equal length with one bearing hairs (arrows; **A, C**). Expelled small ejectisomes had a pointed tip (arrows; **B, F**). From the unfixed specimen, an elongated tubular structure with a globular end was observed (arrows; **D, E**). Scale bars = 5 μm.

**FIGURE 8.**
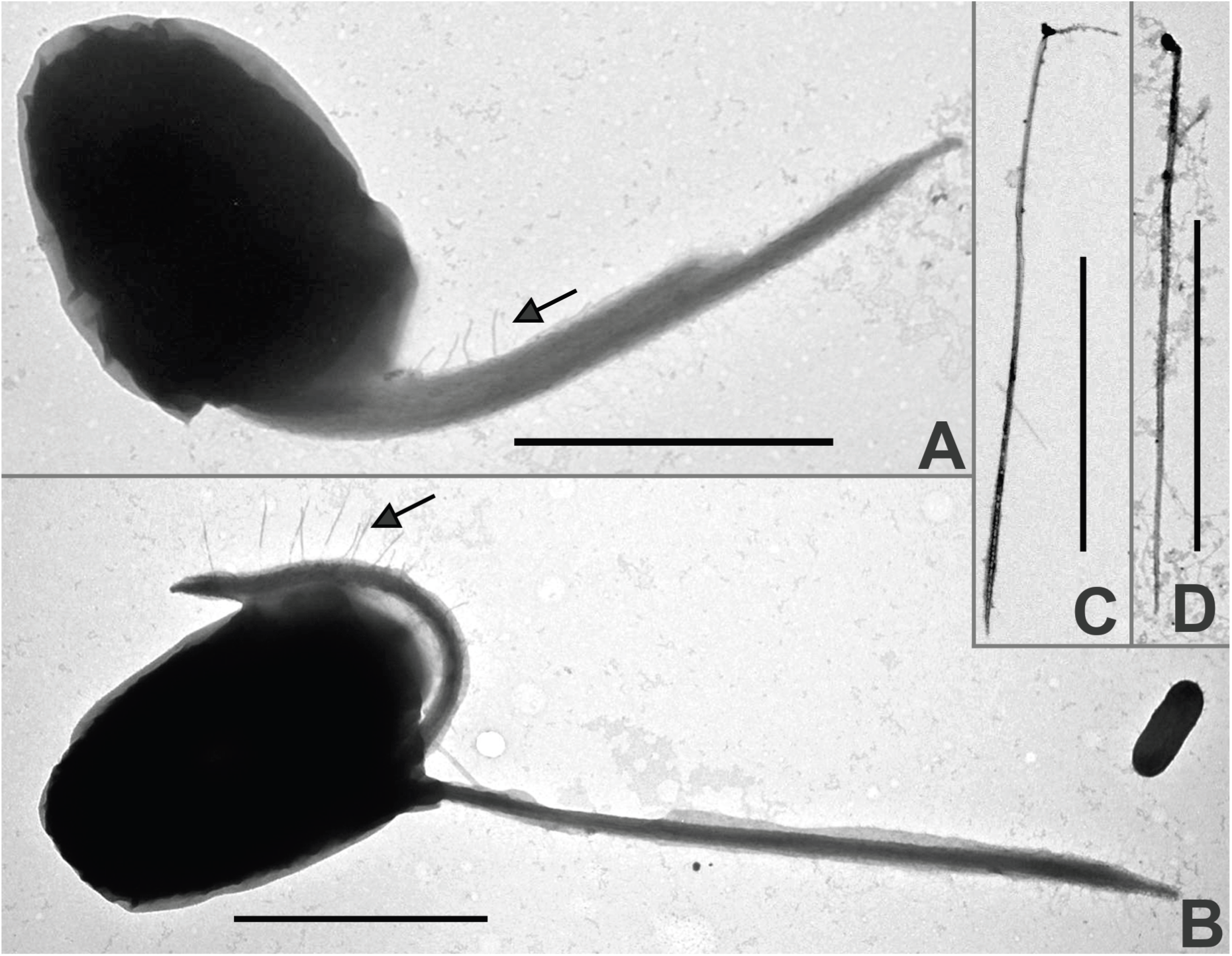
Whole-mount transmission electron microscope (TEM) images from *Goniomonas duplex* strain PT_04. A cell had two unequal flagella with the shorter one bearing hairs (arrows; **A, B**). The small ejectisome was typically longer than the cell length with a pointed tip when expelled from the cell (**C, D**). Scale bars = 5 μm.

The same three strains were also subjected for the standard TEM observation (Figures 9-11). The three were similar in that the nucleus was located towards the dorsal surface where the flagella were inserted (Figures 9A, 10A, 11A) and the mitochondrion had the flat cristate (Figures 9J, 10G, 11G). In addition, the food vacuoles were observed for all three (Figures 9D, 10E, 11A, 11D). Most notably, the area between the nuclear envelope and the rough endoplasmic reticulum was expanded; within this space, fibrous material and small darkly stained globular particles of 130-190 nm in diameter were observed (Figures 9E, 9I, 10C, 11H). In *Goniomonas duplex* PT_04, the vertical sections through the anterior end revealed the opening along the ventral surface (Figure 11E, 11F), suggesting the presence of vertical opening that is continuous to the anterior lateral opening.

**FIGURE 9.**
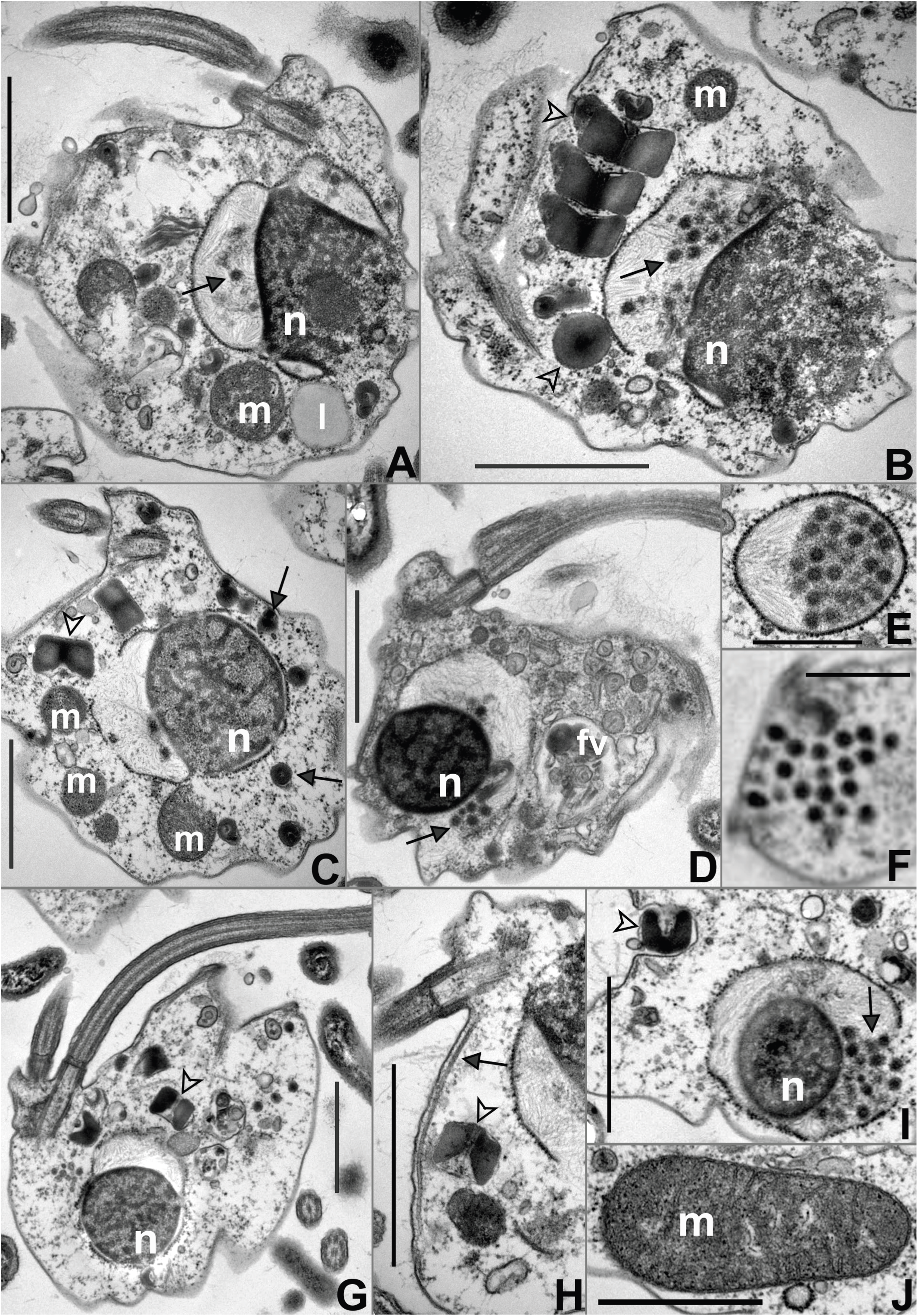
Transmission electron microscope (TEM) images for *Goniomonas ullengensis* strain UH_01. Longitudinal (**A, D, G**) and transverse (**B, C**) sections showed the nucleus(n), a mitochondrion (m), a food vacuole (fv), and a lipid body (l). The rough endoplasmic reticulum was notably enlarged around the nucleus (**A, B, C, D, E, I**); within which thread-like structures and small circular particles were located. The latter particles were also found in an aggregate within the cytoplasm (**F**). The large ejectisome (arrowheads; **B, C, G, H, I**), as well as the small ejectisome (arrows, **C**), had the butterfly shape from the side and the circular shape from the top (or bottom). A microtubular root ran along the anterior cell margin (an arrow; **H**). The mitochondrion had the flat cristae (**J**). Scale bars = 2 µm (1µm for I, J).

**FIGURE 10.**
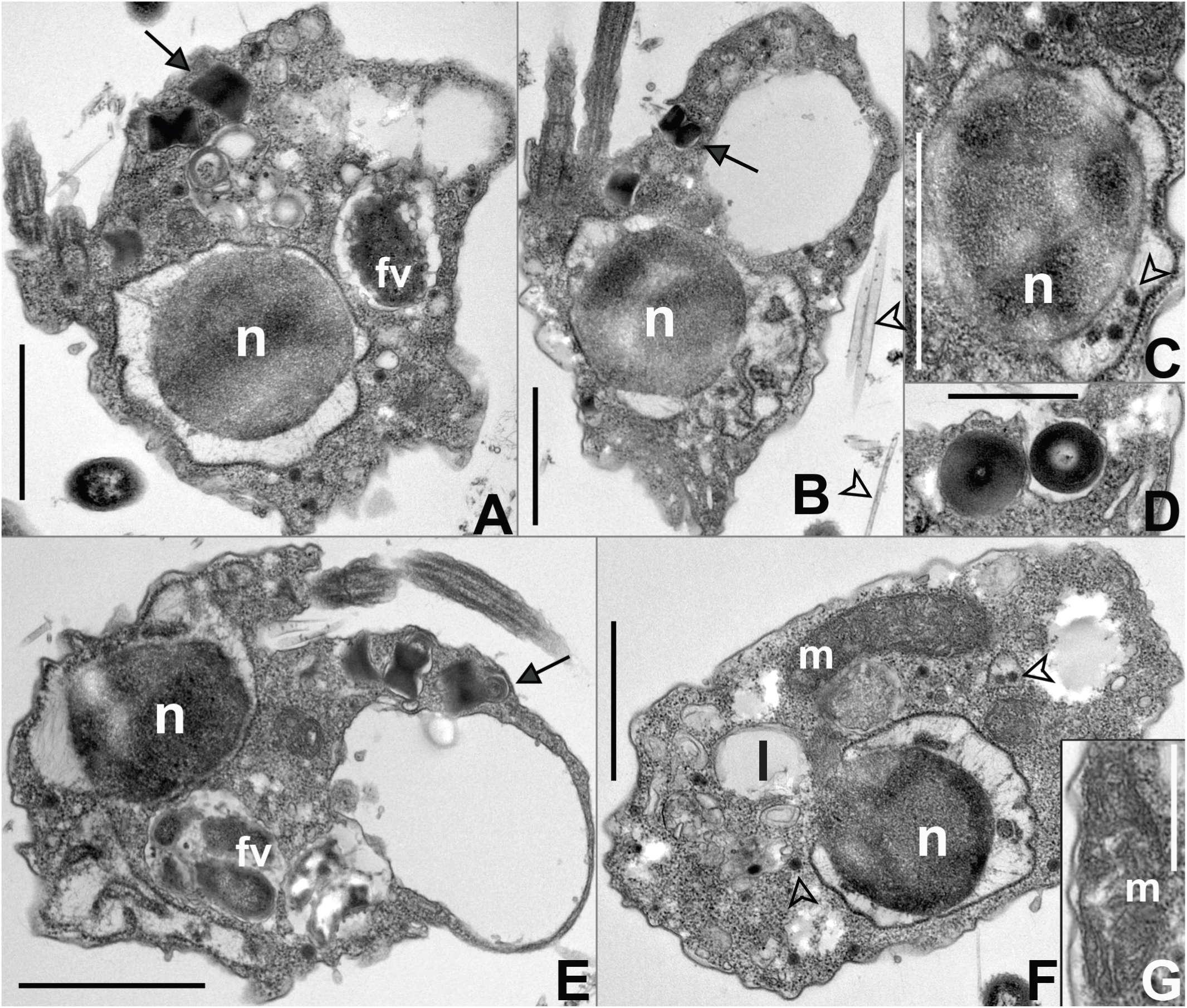
Transmission electron microscope (TEM) images for *Goniomonas lingua* strain GW. The nucleus was located below the area where the flagella emerge (**A, B, E**) and was associated with the rough endoplasmic reticulum (RER). Thread-like structures as well as small dark colored particles were found between the nuclear membrane and RER (arrowhead; **C**). The small particles were also found within the cytoplasm (an arrowhead, **F**). The small and large ejectisome had the butterfly shape from the side and the circular shape from the top (or bottom) (arrows, **A, B, D, E**). An oblique section through the cell revealed an anterior pocket where prey engulfment presumably takes place (**A, B, E**). Food vacuoles were in the vicinity of this large, enclosed hollow (**A, C**). The mitochondrion had the flat cristae (**G**). Abbreviations: fv = food vacuole; l = lipid body; m = mitochondrion; n = nucleus. Scale bars = 2 µm

**FIGURE 11.**
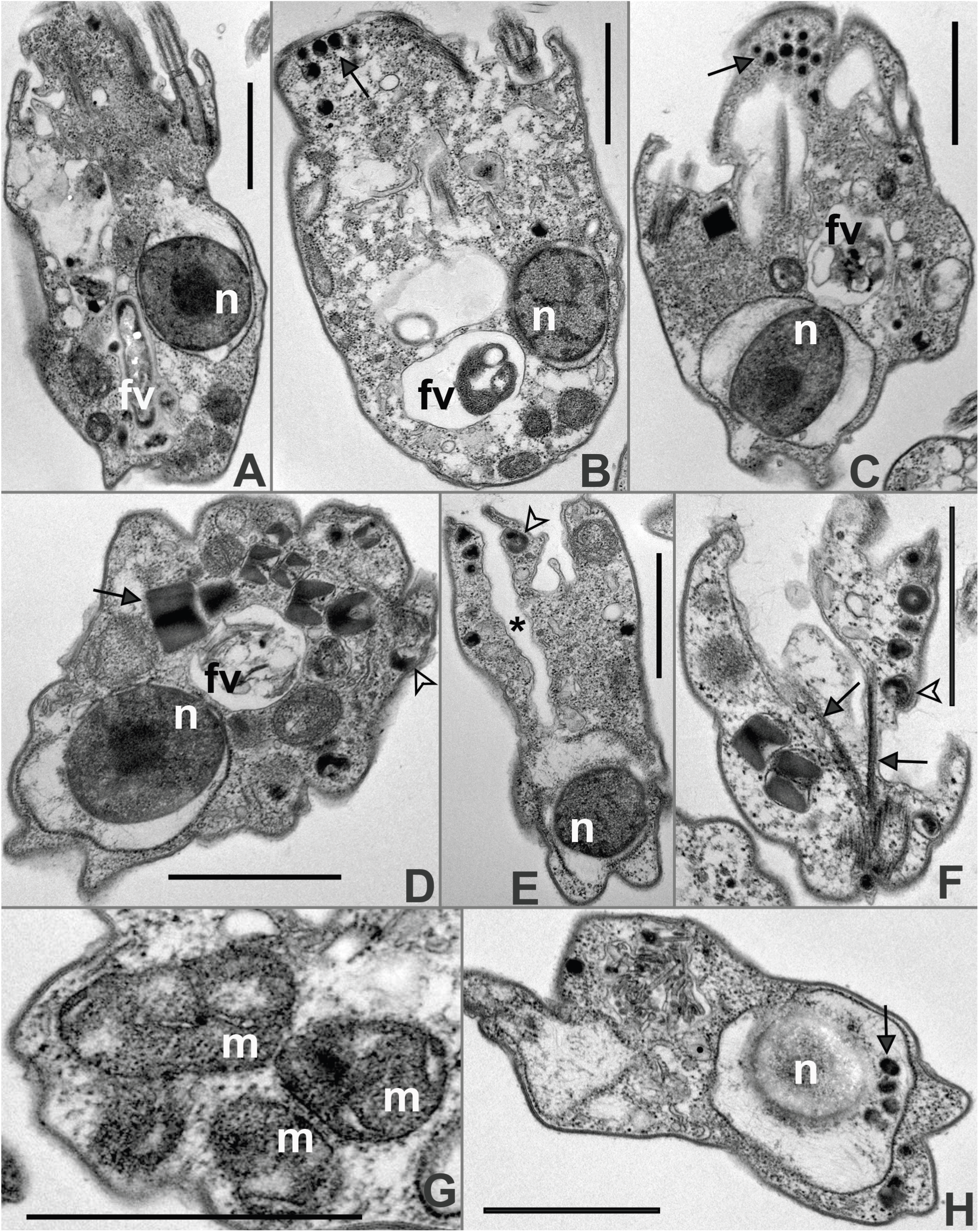
Transmission electron microscope (TEM) images for *Goniomonas duplex* strain PT_04. Longitudinal (**A**) and transverse (**D, E, F**) views of the flagellate. The nucleus was located along the side where the flagellar emerged (**A, B**) and was closely associated with the rough endoplasmic reticulum (RER) (**A, B, C, D, E, H**). The area between the nuclear envelope and RER was enlarged and occupied by thread-like structures and small dark stained particles (arrow, **H**). An aggregate of small particles was found around the cell’s anterior (arrows; **B, C**). The small (arrowhead; **D, E, F**) and large (arrow; **D**) ejectisomes were positioned underneath the cell membrane. Cross sections through the cell anterior (**E, F**) revealed a space between the right and left sides (asterisk symbol, **E**) where the feeding takes place. That opening (“cell’s mouth”) was supported by microtubular roots (arrows, **F**). The mitochondrion had the flat cristae (**G**). Abbreviations: fv = food vacuole; m = mitochondrion; n = nucleus. Scale bars = 2 µm

### Molecular sequence analyses

Phylogenetic analyses of the 18S rDNA sequences placed the newly established strains within the marine clade of the Goniomonadophyceae, which was clearly separated from the freshwater clade comprising long-branching taxa (Figure 12). Within the marine clade, our strains fell into four distinct sub-lineages with one branching very closely with *Goniomonas avonlea* CCMP3327. Our ten strains, including UH_01, were clustering together with the strain MBIC11052; we suggest this sub-lineage represents *G. ulleungensis*. Four of our strains, including GW, formed a separate sub-lineage alongside three previously established strains by other researchers; we recognize this sub-lineage as *G. lingua*. The rest were forming a separate sub-lineage, but given the genetic variation seen among them and absence of morphological data for most, the only small branch including PT_04 was assigned as *G. duplex*. The *G. avonlea* branch was sister to the rest of marine strains; this topology was well supported by both maximum likelihood and maximum parsimony analyses. *G. ulleungensis* and *G. lingua* were closely related to each other to the exclusion of other marine strains.

**FIGURE 12.**
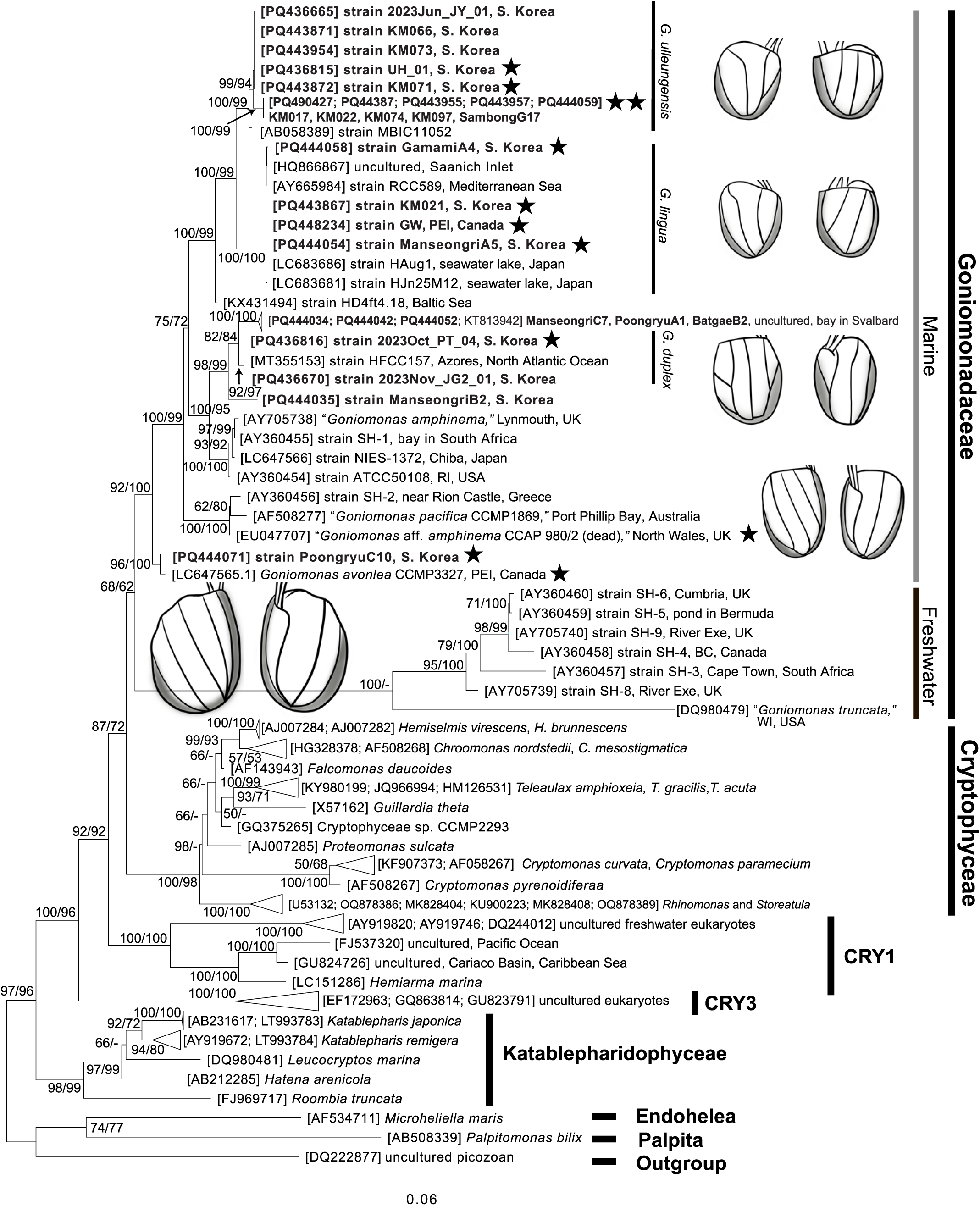
Maximum likelihood (ML) tree based on the analyses of 18S rDNA sequences. ML and maximum parsimony bootstrap values of 50% and greater are shown at the corresponding nodes. The newly acquired sequences in the study are shown in bold face. The scale bar represents the nucleotide substitutions per site. Those strains for which the morphological data are available are denoted with an asterisk symbol.

Pairwise 18S rDNA sequence analyses of marine goniomonad sequences revealed considerable genetic distances among the marine sub-lineages (Table 1). The percent difference between *Goniomonas ulleungensis* and *G. lingua* was 2.8% whereas *G. duplex* differed from either of these by more than 6%. *G. avonlea* differed from any other marine goniomonads by 5.4% or greater.

### Taxonomic summary

Eukaryota: Cryptista Adl et al., 2019; Phylum Cryptophyta Cavalier-Smith, 1986; Class Goniomonadophyceae Cavalier-Smith, 1993 (a typified name; Guiry, 2024); Genus *Goniomonas* Stein, 1878.

#### Goniomonas

##### Diagnosis

Colorless unicellular protist with two flagella that emerge from a corner of the anterior end. Cells flattened with the rounded (to somewhat angular) posterior and the truncated or obtuse anterior. A conspicuous row of the large ejectisomes along the anterior end. Cells covered with rectangular periplast plates that are vertically arranged. Numerous smaller ejectisomes localized underneath the cell membrane in between the periplast plates. Cells skidding on the surface and swimming in the water column. Mitochondrion with the flat cristae. Plastids absent. Bacterivorous. Found in both freshwater and marine habitats.

##### Type species

*Goniomonas truncata* (Fresenius 1858) Stein 1878

#### *Goniomonas ulleungensis* sp. nov

##### Diagnosis

Cells 3-5µm long, 3-4µm wide. Two almost equal flagella, slightly longer than the cell length, arise from an anterior corner. When skidding, the cell’s left side faces the surface. The anterior flagellum bears hairs whereas the other oriented laterally is smooth. Three periplast plates are displayed on the left side and four on the right side. The anterior middle portion of the left side is protruded upwards, with the raised side with a rough texture observable from the right side.

##### Hapantotype

One SEM specimen (MABIK FL00030793), deposited at the National Marine Biodiversity Institute of Korea (Seochun-gun, South Korea)

##### Parahapantotype

One TEM bloc (MPRBK-0001a), deposited at the Marine Protist Resource Bank of Korea (MPRBK) located in Ewha Womans University (Seoul, South Korea)

##### DNA sequence

Two 18S rDNA sequences (PQ436815, PQ432910).

##### Type locality

Sandy beach at Hyeonpo Port, Ulleung-do, South Korea (latitude, longitude = 37.5272, 130.8296)

##### Collection date

October 18^th^, 2023.

##### Type strain

The strain UH_01 used for preparing the hapantotype and parahapantotype is maintained at MPRBK as MPRBK-0001. The live culture material can be obtained through the Marine Bio-Resource Information System (www.mbris.kr) or by contacting the correspondence author.

##### Etymology

The specific epithet *ulleungensis* refers to the name of the island (Ulleung-do, or Ulleung island) where the type strain was collected.

##### Zoobank registration

urn:lsid:zoobank.org:act:77C39837-4B8E-48EE-AB68-4F70BF2D247F.

#### *Goniomonas lingua* sp. nov

##### Diagnosis

Cells 3.5-5µm long, 3-4µm wide. Two almost equal flagella, slightly longer than the cell length, arise from an anterior corner. The cell’s left side faces the surface when skidding. The anterior flagellum bears hairs whereas the posterior one is smooth. There are three periplast plates on both the right and left sides. The anterior middle portion of the left side is protruded upwards, with the raised side with its rough interior texture observable from the right side. *Goniomonas lingua* and *G. ulleungesis* are genetically distinguished despite their similar morphology and movement pattern seen under a light microscope. Nonetheless, the two closely related species can be distinguished by the periplast plate pattern; *G. ulleungesis* has one additional periplast plate in the rightmost part on the right side.

##### Hapantotype

One SEM specimen (MABIK FL000307974), deposited at the National Marine Biodiversity Institute of Korea (Seochun-gun, South Korea)

##### Parahapantotype

One TEM bloc (MPRBK-0003a), deposited at the Marine Protist Resource Bank of Korea (MPRBK) located in Ewha Womans University (Seoul, South Korea); One SEM specimen (MABIK FL0003108)—derived from the strain KM021—deposited at the National Marine Biodiversity Institute of Korea (Seochun-gun, South Korea)

##### DNA sequence

Two 18S rDNA sequences (PQ448234, PQ448237).

##### Type locality

Greenwich beach, Prince Edward Island, Canada (latitude, longitude = 46.4503, - 62.7125)

##### Collection date

August 27^th^, 2011.

##### Type strain

The strain GW used for preparing the hapantotype and one parahapantotype is maintained at MPRBK as MPRBK-0003. The culture material can be obtained through the Marine Bio-Resource Information System (www.mbris.kr) or by contacting the correspondence author.

##### Etymology

The specific epithet *lingua*, tongue, refers to the (tongue-like) anterior extension from the left side. Given its proximity to the cell’s mouth, the structure may be used for prey recognition.

##### Zoobank registration

urn:lsid:zoobank.org:act:C88EA624-F1F3-4D01-95D7-11EEDAB6B9B9;

#### *Goniomonas duplex* sp. nov

##### Diagnosis

Cells 4.5-5.5µm long, ∼4µm wide. Two unequal flagella arise from an anterior corner. The shorter anterior flagellum, a slightly longer than the cell length, bears hairs whereas the longer posterior flagellum, 1.5 times the cell length, is smooth. The cell’s right-side faces towards the surface when skidding. There are five periplast plates on the left side and three on the right side. The leftmost periplast area in the left side is horizontally split into two pieces.

##### Hapantotype

One SEM specimen (MABIK FL00030795), deposited at the National Marine Biodiversity Institute of Korea (Seochun-gun, South Korea)

##### Paratype

One TEM bloc (MPRBK-0002a), deposited at the Marine Protist Resource Bank of Korea (MPRBK) located in Ewha Womans University (Seoul, South Korea).

##### DNA sequence

Two 18S rDNA sequences (PQ436816, PQ436667).

##### Type locality

Padori beach, Taean-gun, Chungcheongnam-do, South Korea (latitude, longitude = 36.7440, 126.1324)

##### Collection date

October 31^st^, 2023.

##### Type strain

The strain PT_04 used for preparing the hapantotype and parahapantotype is maintained at MPRBK as MPRBK-0002. The culture material can be obtained through the Marine Bio-Resource Information System (www.mbris.kr) or by contacting the correspondence author.

##### Etymology

The specific epithet *duplex*, twofold, refers to the left periplast region of the left side that is divided into two compartments.

##### Zoobank registration

urn:lsid:zoobank.org:act:104A4B6B-366C-471E-82C2-98B2A1DEACC4

## DISCUSSION

In this study, we describe three new marine goniomonad species by light microscopy, electron microscopy, and 18S rDNA gene phylogenetic inference. Our molecular phylogenetic analyses placed all the newly acquired marine strains, 21 in total, into the marine goniomonad clade (Figure 12). In agreement with previous studies (e.g. von der Heyden et al., 2004), our results based on the updated sequence matrix support a deep divergence between the marine and freshwater goniomonads, with the latters being farily long-branching. Within the marine clade, several genetically distinct sub-lineages were identified with the clade comprising *Goniomonas avonlea* being sister to the rest of marine species. Despite its widespread distribution in aquatic environments (Lee, 2002), *Goniomonas* has been neglected for in-depth descriptive studies. For example, with exceptions for *G. avonlea* CCMP3327 (Kim and Archibald, 2013) and *G.* aff. *amphinema* (Martin-Cereceda et al., 2010), all the previously acquired marine goniomonad 18S rDNA sequences do not have matching morphological data. Our study fills in this gap, at least partially, by acquiring and comparatively analyzing both 18S rDNA and morphological data from new marine isolates from South Korea and Canada.

Goniomonads are genetically diversified in 18S rDNA (Figure 12; von der Heyden et al., 2004), but their light microscopic features appear well conserved. *Goniomonas* species share morphological features, including the flattened, colorless, nano-sized body, presence of a conspicuous transverse band along the truncated/obtuse anterior end, and two flagellar inserted at an anterior depression (Larsen and Patterson, 1990). This gross morphological conservation has contributed to the taxonomic under-splitting of the genus *Goniomonas* (von der Heyden et al., 2004) but our careful morphological characterization using a combination of light and electron microscopic techniques revealed that distinct genotypes (or sub-lineages) of marine goniomonads can be differentiated by morphological data as well.

To date, including this study, five marine goniomonad species have been characterized using both morphological and 18S rDNA data. These include *Goniomonas* aff. *amphinema* (Martin-Cereceda et al., 2010), *G. avonlea* (Kim and Archibald, 2013), *G. ulleungensis, G. lingua*, and *G. duplex*. *Goniomonas* aff. *amphinema* was named so as it closely resembles *G. amphinema* as originally described by Larson and Patterson (1990) with a few exceptions. Martin-Cereda et al. (2010) noted that *G*. aff. *amphinema* has the anterior flagellum that is slightly shorter than the cell, whereas *G. amphinema* was originally described to have the anterior flagellum about half the cell length. In addition, *G*. aff. *amphinema* has the average width of 3.5 µm (Martin-Cereceda et al., 2010), which is slightly narrower than the original description for *G. amphinema* (i.e. 4-6 µm in width; Larson and Patterson, 1990). *G. duplex* is also similar to *G. amphinema* (as described by Larson and Patterson, (1990)), but it has the anterior flagellum that is slightly longer than the cell (Figure 1J, 1L), whereas *G. amphinema* has a much shorter anterior flagellum. This species is distinguished from *G.* aff. *amphinema* by 5.9% difference in 18S rDNA (Table 2). Morphologically, *G. duplex* and *G*. aff. *amphinema* are quite similar in size and flagellar orientation, but the flagella of *G. duplex* are slightly longer. Furthermore, their periplast plate pattern on the left side differs in that *G. duplex* has its leftmost periplast area laterally split into two pieces (Figure 5B, 5E), which is not the case for *G*. aff. *amphinema*. *Goniomonas avonlea* is similar to *G.* aff. *amphinema* and *G. duplex* in terms of flagellar orientation with the posterior flagellum trailing posteriorly. The periplast pattern is very similar between *G. avonlea* and *G.* aff. *amphinema*: three periplast strips on the right side and four on the left side. Nonetheless, *G. avonlea* can be easily distinguished from *G.* aff. *amphinema* or *G. duplex* by its larger cell body (Figure 2). Also note that *G. avonlea* differs from four other described marine goniomonad species in 18S rDNA by more than 5% (Table 2).

**Table 2.** Pairwise comparison of seven marine *Goniomonas* 18S rRNA gene sequences. The values shown are pairwise sequence percent identities across the 18S rRNA gene region that corresponds to the positions 90 and 1,716 within the *Chlamydomonas reinhardtii* sequence M32703.

|  | PQ436815 | PQ448234 | PQ436816 | AY705738 | AF508277 | EU047707 |
| --- | --- | --- | --- | --- | --- | --- |
| *G. ulleungensis*<br>UH_01 (MPRBK-0001) [PQ436815] |  |  |  |  |  |  |
| *G. lingua*<br>GW (MPRBK-0002) [PQ448234] | 97.2% |  |  |  |  |  |
| *G. duplex*<br>PT_04 (MPRBK-0003) [PQ436816] | 93.3% | 93.2% |  |  |  |  |
| "*G. amphinema*" [AY705738] | 93.2% | 93.4% | 96.2% |  |  |  |
| "*G. pacifica* CCMP1869"<br>[AF508277] | 91.4% | 91.4% | 92.0% | 91.8% |  |  |
| "*G. aff. amphinema*" [EU047707] | 93.9% | 94.0% | 94.1% | 93.7% | 97.2% |  |
| *G. avonlea* CCMP3327 [JQ434475] | 93.5% | 93.4% | 94.1% | 94.3% | 92.0% | 94.6% |

*Goniomonas ulleungensis* and *G. lingua* can be easily differentiated from other three marine goniomonad species by having two equal to subequal flagella orienting anterior-laterally. These two species also share a characteristic tongue-like protrusion that is visible from the right side (Figures 3A, 4A). Rather unexpectedly, the cell orientation for these speices during skidding was found to be the opposite of that for other three species. While the skidding cells of *G. avonlea* and *G. duplex* (and likely *G*. aff. *amphinema*) have its right side facing the surface, *G. ulleungensis* and *G. lingua* have its left side orienting towards the surface. Given the phylogenetic relationships of the five species, the most parsimonious scenario is that the common ancestor of marine goniomonads had its right side facing the surface while skidding with one flagellum trailing and the other directed anteriorly. The direction of the skidding cells flipped and one flagellum changed its orientation from the posterior to the anterior-lateral direction at the ancestry for the *G. ulleungensis-G. lingua* clade. While *G. ulleungensis* and *G. lingua* are difficult to differentiate by light microscopy, they differ by 2.8% in 18S rDNA region and also differ in ultrastructure. Specifically, *G. ulleungensis* has four periplast plates on the right side whereas *G. lingua* has three (Figures 3F, 4A, S2C).

In characterizing the goniomonad morphology, we note that, variation in the cell length and width was observed even among strains from the same sub-lineage (as defined here by having >99% identity in 18S rDNA region; Figure 2). For example, *Goniomonas ulleungensis* KM071 is markedly smaller than other three strains, and it is slightly stretched out laterally, whereas the others are vertically elongated. Such size variation among strains of the same genotype could be explained by different growth conditions (e.g., growth media, temperature), growth phase, or sexual dimorphism (as noted for some plastid-bearing cryptophyte species; Altenburger et al., 2020). Even so, two strains of *G. avonlea* for which we have size measurement data were notably larger than all other surveyed marine goniomonad strains. This supports the utility, albeit limited, of cell size data in marine goniomonad identification.

Based on analyses of existing sequence data from isolates or environmental samples (e.g. Weber et al., 2017), it is apparent that more marine goniomonad species remain to be explored and described. For future taxonomic study of goniomonads, we recommend that researchers acquire 18S rDNA data alongside the following morphological data. From light microscopic observation of living cells, the cell size, flagellar length/orientation as well as the orientation of the skidding cell can be determined. We note that while some researchers suggested the utility of the cell length-to-width ratio in delineating cryptophyte species (Klaveness, 1985), we did not find this feature particularly helpful for marine goniomonad taxonomy (Figure 2). The whole-mount TEM preparation can be done relatively quickly compared to the standard TEM or SEM work, and we recommend this method over SEM for characterizing the flagellar appendages and the discharged ejectisomes. We note that SEM images can be misleading at times especially if the appendages are damaged or appressed firmly to the flagellum during the specimen preparation. For species-level delineation by morphology, characterization of the periplast plate pattern by SEM is especially helpful. The standard TEM data did not reveal any obvious differences between the marine goniomonad species although the flagellar apparatus characterization might reveal additional ultrastructural features suitable for species-level delineation of gonimonads. Even so, TEM data revealed an interesting commonality shared among the three marine goniomonads (but not *Goniomonas avonlea*; Kim and Archibald, 2013) as well as one freshwater goniomonad strain investigated by Mignot (1965). These four goniomonads displayed an area, perhaps as large as the volume of the nucleus, located between the nucleus and the rough endoplasmic reticulum where fibrous material and small spherical particles reside (Figures 9A, 9B, 10C, 11H; Planche IV, Fig. b in Mignot (1965)). Mignot (1965) determined that the small spherical inclusions are glycogenic in nature. These might be energy storage molecules or precursors to ejectisomes; further studies are needed to understand the functional aspects of these structures.

## Supporting information

DATA_S1_SSUaln

FigS1_LMs

FigS2_SEM

## ACKNOWLEDGEMENTS

This work was supported by the National Marine Biodiversity Institute of Korea (MABIK): 1) the 2024 project Management of the Marine Fishery Bio-resources Center and 2) the project ID 2024M00200.We thank Dr. Michelle Leger for her help with sampling at PEI, Canada; Jaung-Mi Han and Mi-Jin Ji at the Nano Imaging Laboratory in the Ewha Medical Research Institute for their help with transmission electron microscopy.

