## Supplementary figures and images for "Morphological and Molecular Phylogenetic Characterization of Three New Marine Goniomonad Species"

### FigS1_LMs

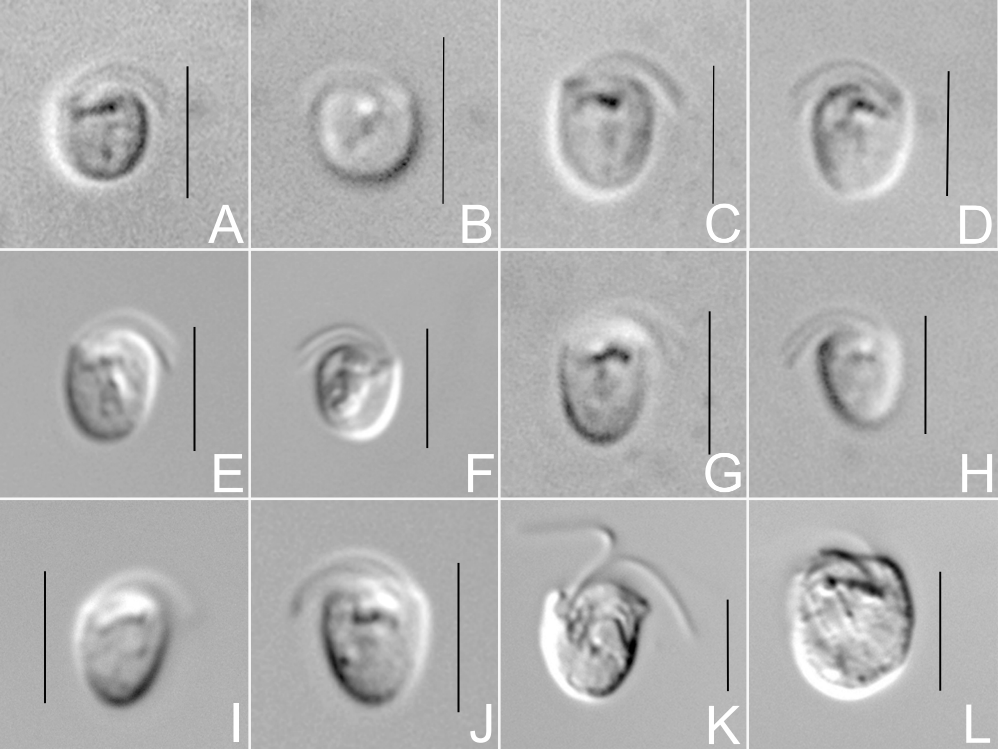

### FigS2_SEM

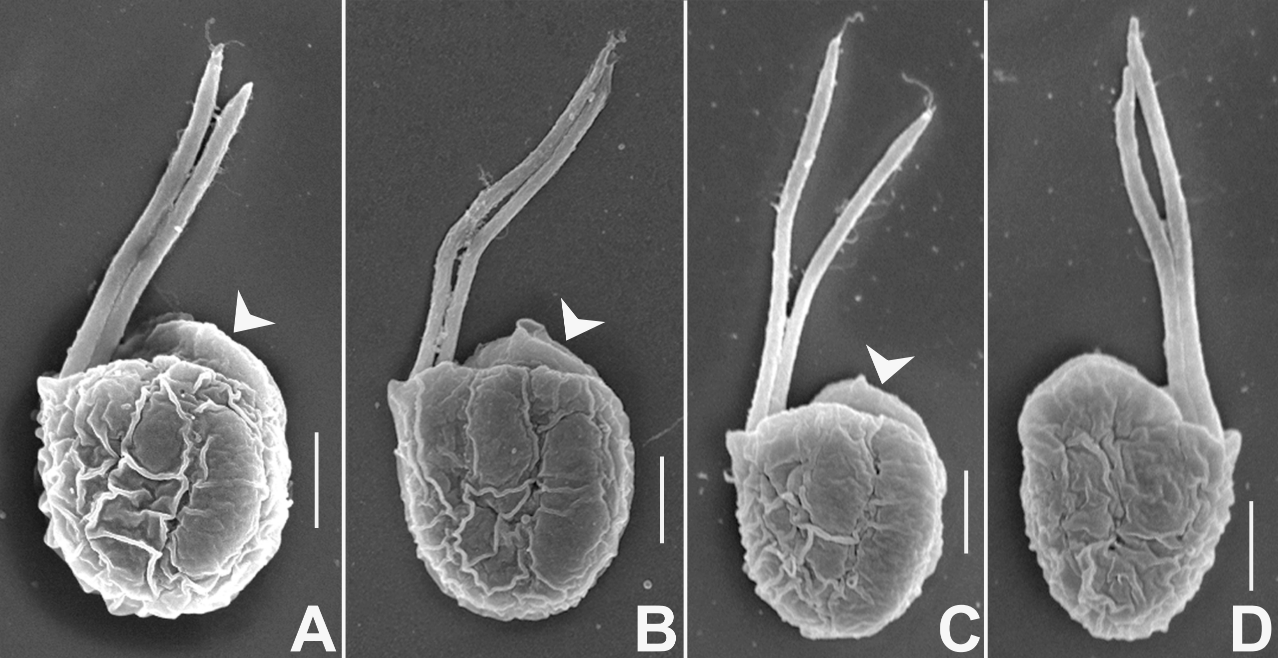
