## Supporting_Information for "Morphological and Molecular Phylogenetic Characterization of Three New Marine Goniomonad Species"

1. Table S1: PCR primers used to amplify 18S rDNA regions in this study. Primer positions are relative within the *Chlamydomonas reinhardtii* sequence M32703
2. Figure S1: Light microscope images of additional marine goniomonad strains newly obtained in this study. **A-F**. *Goniomonas ulleungensis* strains KM071 (A, B), KM097 (C, D), SambongG17 (E, F) **G-J**. *Goniomonas lingua* strains KM021 (G, H), ManseongriA5 (I), GamamiA4 (J). **K-L**. *Goniomonas avonlea* strain PoongryuC10. Scale bars = 5 μm
3. Figure S2: Scanning electron microscope (SEM) images for *Goniomonas ulleungensis* strains KM071 (**A**) & KM097 (**B**), and *G. lingua* strain KM021 (**C, D**). From the right side view, an anterior protrusion from the left side of the cell (arrowheads in A, B, C) is visible. Scale bars = 1 µm
4. Data S1: 18S rDNA sequence alignment used in this study (nexus format)
5. Data S2: Cell size measurement data
6. Movie S1: A video showing the movement of *Goniomonas ulleungensis* strain UH_01
7. Movie S2: A video showing the movement of *Goniomonas lingua* strain GW
8. Movie S3: A video showing the movement of *Goniomonas duplex* strain PT_04

Table S1. PCR primers used to amplify 18S rDNA regions in this study. Primer positions are relative within the *Chlamydomonas reinhardtii* sequence M32703.

| Primer name | | Relative positions (5′ to 3′) | Sequence (5′ to 3′) | Reference |
| --- | --- | --- | --- | --- |
| Pair1 | nu-SSU-0024-5′ | 0004–0024 | CTG GTT GAT CCT GCC AGT AGT | Kim et al. 2006 |
|  | nu-SSU-1757-3′ | 1777–1757 | CAG GTT CAC CTA CGG AAA CCT | Kim et al. 2006 |
| Pair2 | E572F | 0570–0587 | CYG CGG TAA TTC CAG CTC | Comeau et al. 2011 |
|  | E1009R | 1005–0986 | AYG GTA TCT RAT CRT CTT YG | Comeau et al. 2011 |
| Pair3 | 18S_CrN1F | 0014–0037 | CTG CCA GTA GTC ATA TGC TTG TCT | Choi et al. 2013 |
|  | 18S_BRK | 1764–1764 | GGA AAC CTT GTT ACG ACT TCT C | Choi et al. 2013 |
| Pair4 | EukA | 0001-0021 | AAC CTG GTT GAT CCT GCC AGT | Medlin et al. 1988 |
|  | EukB | 1788-1765 | TGA TCC TTC TGC AGG TTC ACC TAC | Medlin et al. 1988 |
